# The CD11a and EPCR marker combination simplifies and improves the purification of mouse hematopoietic stem cells

**DOI:** 10.1101/219063

**Authors:** Alborz Karimzadeh, Vanessa Scarfone, Connie Chao, Karin Grathwohl, John W. Fathman, David Fruman, Thomas Serwold, Matthew A. Inlay

## Abstract

Hematopoietic stem cells (HSCs) are the self-renewing multipotent progenitors to all blood cell types. Identification and isolation of HSCs for study has depended on the expression of combinations of surface markers on HSCs that reliably distinguish it from other cell types. However, the increasing number of markers required to isolate HSCs has made it tedious, expensive, and difficult for newcomers, suggesting the need for a simpler panel of HSC markers. We previously showed that phenotypic HSCs could be separated based on expression of CD11a, and that only the CD11a negative fraction contained true HSCs. Here, we show that CD11a and another HSC marker, EPCR, can be used to effectively identify and purify HSCs. We introduce a new two-color HSC sorting method that can highly enrich for HSCs with efficiencies comparable to the gold standard combination of CD150 and CD48. Our results demonstrate that adding CD11a and EPCR to the HSC biologist’s toolkit improves the purity of and simplifies isolation of HSCs.

**Significance Statement:** The study of hematopoietic stem cells (HSCs) and their purification for transplantation requires a panel of surface markers that can be used to distinguish HSCs from other cell types. The number of markers necessary to identify HSCs continues to grow, making it increasingly difficult to identify HSCs by flow cytometry. In this study, we identified a combination of two surface markers, CD11a and EPCR, to enrich for HSCs in the mouse bone marrow without the need for additional markers. This simplified panel could aid HSC research by reducing the number of markers necessary to identify and isolate HSCs.

## Introduction

Hematopoietic stem cells (HSCs) are the self-renewing, multipotent, and engraftable cells of the blood system [1]. Successful HSC transplantation (HSCT) can potentially treat any disorder inherent to the hematopoietic system by ablation of the defective blood system followed by reconstitution by healthy donor HSCs [2]. However, HSCT is reserved only for high-risk patients due to the dangers of HSCT-related complications, including graft rejection, graft failure, and graft versus host disease [2]. Transplantation of sufficient numbers of pure HSCs can bypass many of these HSCT-related complications [3-5]. Therefore, much effort has been invested in strategies to improve the purity of donor HSCs.

HSCs are identified by their expression of a combination of molecules on their cell surface called surface markers. In mice, the ever-growing list of surface markers whose positive or negative expression marks HSCs includes CD34, Kit, Sca-1, Lineage (a cocktail of markers of mature lineages), CD27, CD48, CD150, FLK2, CD9, EPCR and others [6-9]. The marker combination Kit+ Lineage- Sca-1+, which defines the “KLS” population (also called “LSK” or “KSL”), contains all HSCs and multipotent progenitors in the bone marrow (BM). To isolate long-term HSCs within the KLS population, additional marker combinations are needed such as 1) Flk2+ CD34-, 2) CD48- CD150+, or 3) CD150+ CD34- [6, 10, 11]. However, the increasing number of markers needed to purify HSCs (currently around 6-8), the nuances of each of the fluorochromes and antibodies required for optimal staining and gating, and the long and expensive assays needed for gating validation have made it difficult for newcomers to properly identify and sort HSCs. Furthermore, many of these markers can change expression during stressful conditions such as upon inflammation or after irradiation, making many of them unreliable for identifying HSCs in these contexts [12-14]. Therefore, there remains a need for simpler and more inclusive strategies for marking and identifying HSCs in healthy and challenged BM.

We previously introduced CD11a as a new marker to isolate HSCs. CD11a (integrin alpha L, or *Itgal*) heterodimerizes with the β2 integrin CD18 to form LFA1. LFA1 interacts with ICAM-1 and has roles in transendothelial migration, activation, and differentiation of lymphocytes [15-18]. We found that while CD11a is expressed on nearly all hematopoietic lineages, it is downregulated in HSCs [19]. We showed that stringently-gated adult HSCs can be separated into CD11a+ and CD11a- fractions, with only the CD11a- fraction showing long-term engraftment upon transplantation. This was not due to antibody binding to the CD11a+ cells (potentially blocking LFA1-mediated migration), as inclusion of the CD11a antibody affected neither BM homing, nor long-term engraftment of HSCs. These findings suggested that CD11a should be added to the marker panel when isolating HSCs at the highest level of purity. Here, we introduce an alternative strategy for identification and sorting of HSCs with the use of CD11a and EPCR (endothelial protein C receptor, *Procr*, CD201) as another efficient HSC marker. We show that CD11a and EPCR can be used with classical HSC markers to purify HSCs, but furthermore, can be used alone as a simple two-color method to highly enrich for HSCs.

## Materials & Methods

### Mice

C57Bl/6 (stock no. 00664), CK6/ECFP (TM5; stock no. 004218), and mT/mG (stock no. 007576) strains from Jackson Laboratory (Bar Harbor, ME, USA) were utilized as donors/recipients/helpers. All strains were maintained at the Gross Hall and Med Sci A vivarium facilities at UCI and fed with standard chow and water. All animal procedures were approved by the International Animal Care and Use Committee (IACUC) and University Laboratory Animal Resources (ULAR) of University of California, Irvine.

### Antibodies

For list of antibodies, refer to Table S1 (“Antibodies Table”) in Supporting Information.

### Cell sorting

For flow cytometry, BM was harvested from tibias and femurs by flushing with ice-cold FACS buffer (PBS + 2% fetal bovine serum) followed by red blood cell lysis by ACK lysing buffer and filtration through a 70 µ mesh. BM was harvested from donor mice by crushing leg bones in ice-cold FACS buffer followed by red blood cells lysis by ACK lysing buffer and filtration through a 70 µ mesh to remove debris. Where indicated, BM was Kit enriched using anti-Kit (anti-CD117) microbeads on an AutoMACS (Miltenyl Biotec). Cells were stained with antibodies listed in Table S1 (“Antibodies Table”) in FACS buffer. Cells were sorted on a BD FACS-Aria II (Becton Dickinson) into ice-cold FACS buffer for transplantation.

### Transplantation, and blood and BM analysis

Defined numbers of HSCs (as indicated in each experiment) were transplanted by retro-orbital injection into lethally-irradiated isoflurane-anesthetized recipients alongside helper BM from congenically distinguishable C57BL/6 mice. Lethal doses of X-ray irradiation were 800 Rads for single dose, or 950 Rads split dose (XRAD 320, Precision X-ray). Transplanted recipients were fed an antibiotic chow of Trimethoprim Sulfa (Uniprim, Envigo) for 4 weeks post transplantation to prevent potential bacterial infections. For peripheral blood analysis, blood was obtained from the tail vein of transplanted mice at various time points, and red blood cells were depleted using ACK lysis buffer. For BM analysis, BM was harvested from tibias and femurs by flushing with ice-cold FACS buffer followed by ACK lysis and filtration. Cells were stained with lineage antibodies and analyzed on the BD FACS-Aria II. For a comprehensive list of markers used for identification of each population, refer to Table S2 (“Marker definitions of populations analyzed”) in Supporting Information. FlowJo software (Tree Star) was used for data analysis.

### Statistical analysis

Statistical analysis was performed with GraphPad Prism 5 software (La Jolla, CA).

## Results

### CD11a and EPCR in combination with classical HSC markers reveal a distinct population with enriched HSC activity

CD11a and EPCR have each been shown independently to increase HSC purity when used with conventional HSC markers [19-21]. To assess the efficiency of purifying HSCs using CD11a and EPCR together, we first examined their expression in the KLS population, which contains all hematopoietic stem and multipotent progenitor cells and is often referred to as “HSPCs” (Fig. 1). KLS is traditionally defined as Kit+ Lin- Sca-1+, but we substituted CD27 for the Lineage (Lin) cocktail, an expensive combination of markers for mature hematopoietic lineages. CD27 is expressed on HSCs and MPPs, and together with the red blood cell marker Ter119, can be used in place of Lin [14, 22, 23]. Within the KLS population (Ter119- CD27+ Sca-1+ Kit+), we identified two distinct fractions: a CD11a- EPCR+ population and a CD11a+ population (Fig. 1A). We sorted these two populations (from CFP+ donor mice) and transplanted them into lethally-irradiated Bl/6 adult recipients to determine which population contained long-term engraftable HSCs. 500,000 BM cells from Tomato+ mice were co-transplanted as “helper” BM to protect the recipients from hematopoietic failure following irradiation. Recipients were bled and analyzed for donor chimerism in different blood lineages at four-week intervals (**Supporting Information Fig. S1**). Donor chimerism of total blood cells (CD45+) was significantly higher from the CD11a- EPCR+ KLS population than the CD11a+ KLS population, and this difference increased over time (Fig. 1B). Because granulocytes are short-lived, granulocyte chimerism in the peripheral blood is a more accurate indicator of HSC chimerism in the BM compared to total CD45+ blood cells, which includes long-lived lymphocytes that may have come from lymphoid progenitors or multipotent progenitors. The difference in granulocyte chimerism between CD11a- EPCR+ KLS cells and CD11a+ KLS was even more pronounced than total blood chimerism (Fig. 1C). Furthermore, when examining the BM of the recipients, the CD11a- EPCR+ KLS population had higher donor HSPCs (Fig. 1D). As only HSCs are capable of serial transplantation, we next transplanted whole BM from the primary recipients into secondary hosts. Only the BM of recipients of CD11a- EPCR+ KLS cells gave rise to robust donor chimerism in the secondary hosts, indicating that nearly all HSCs are contained in this population (Fig. 1E). Thus, CD11a and EPCR can be used to isolate HSCs within the KLS fraction of BM.

**Figure 1.**
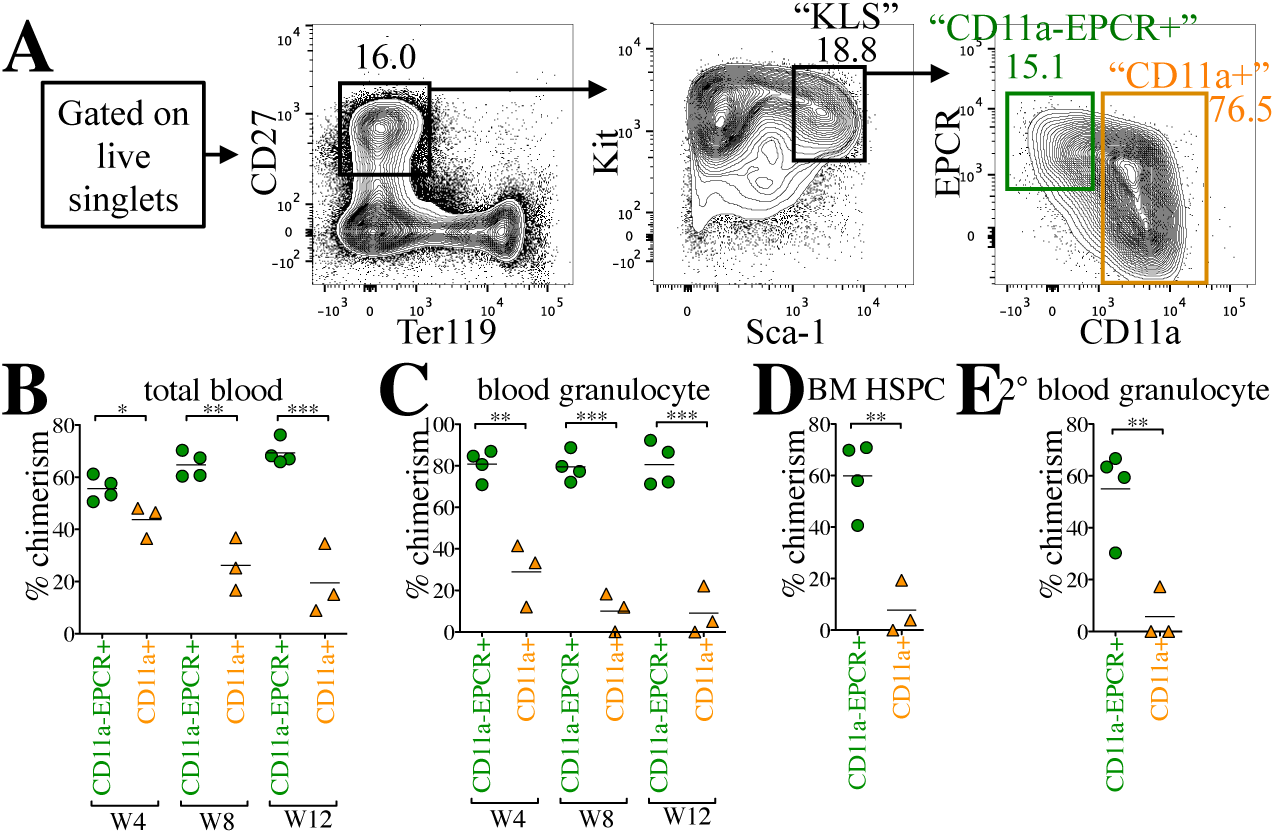
CD11a and EPCR inclusion enriches for HSCs within KLS population. **A)** Representative sorting scheme of CD11a- EPCR+ KLS (green) and CD11a+ KLS (orange) populations from Kit-enriched CFP+ BM. Each sorted population (1,500 CD11a- EPCR+ cells or 10,000 CD11a+ cells per recipient) was transplanted into lethally-irradiated Bl/6 recipients along with 500,000 Tomato+ WBM helper cells. **B-C)** Time-course analysis of donor chimerism in blood. Total (**B**) and granulocyte (**C**) blood chimerism from CD11a- EPCR+ KLS (“CD11a-EPCR+”) and CD11a+ KLS (“CD11a+”) sources in primary recipients at weeks (W) 4, 8, and 12 post-transplant. Total blood was defined as CD45+ and granulocytes as CD45+ Gr1+ Mac-1+. **D)** Donor chimerism of HSPCs in the BM of primary recipients 13 weeks post-transplant. HSPCs are defined as Ter119- CD27+ Sca-1+ Kit+. **E)** Blood granulocyte chimerism in secondary recipients 6 weeks post-secondary transplant. Secondary transplants were done using 1x10^6^ WBM harvested from primary recipients that received “CD11a-EPCR+” or “CD11a+” donor cells. *\*p ≤ 0.05, **p ≤ 0.01, ***p ≤ 0.001 (Student’s unpaired t test).*

### CD11a- EPCR+ KLS directly outcompetes CD11a+ KLS in a competitive transplantation assay

To directly compare HSC activity between the CD11a- EPCR+ and CD11a+ subsets of KLS cells, we performed a competitive transplantation, in which both populations are co-transplanted into the same recipients. In this strategy, recipient mice receive both populations at their physiological ratios, providing a direct comparison of the engraftment efficiency of each. Also, because all KLS cells fall within one fraction or the other, all potential sources of HSCs in the BM are sorted and transplanted. To distinguish the two populations, we sorted one population from CFP-expressing BM, and the other population from Tomato-expressing BM and co-transplanted them along with 100,000 unlabeled Bl/6 helper BM cells (Fig. 2A). In the peripheral blood of the primary recipients, drastically higher percentages of donor total CD45+ cells and granulocytes were derived from the CD11a- EPCR+ KLS compared to the CD11a+ KLS source (Fig. 2B-C). Higher donor chimerism of HSCs in the BM compartment of primary recipients as well as blood granulocyte chimerism in secondary recipients also originated only from the CD11a- EPCR+ KLS donor source, indicating that this population contained all the HSCs (Fig. 2D-E).

**Figure 2.**
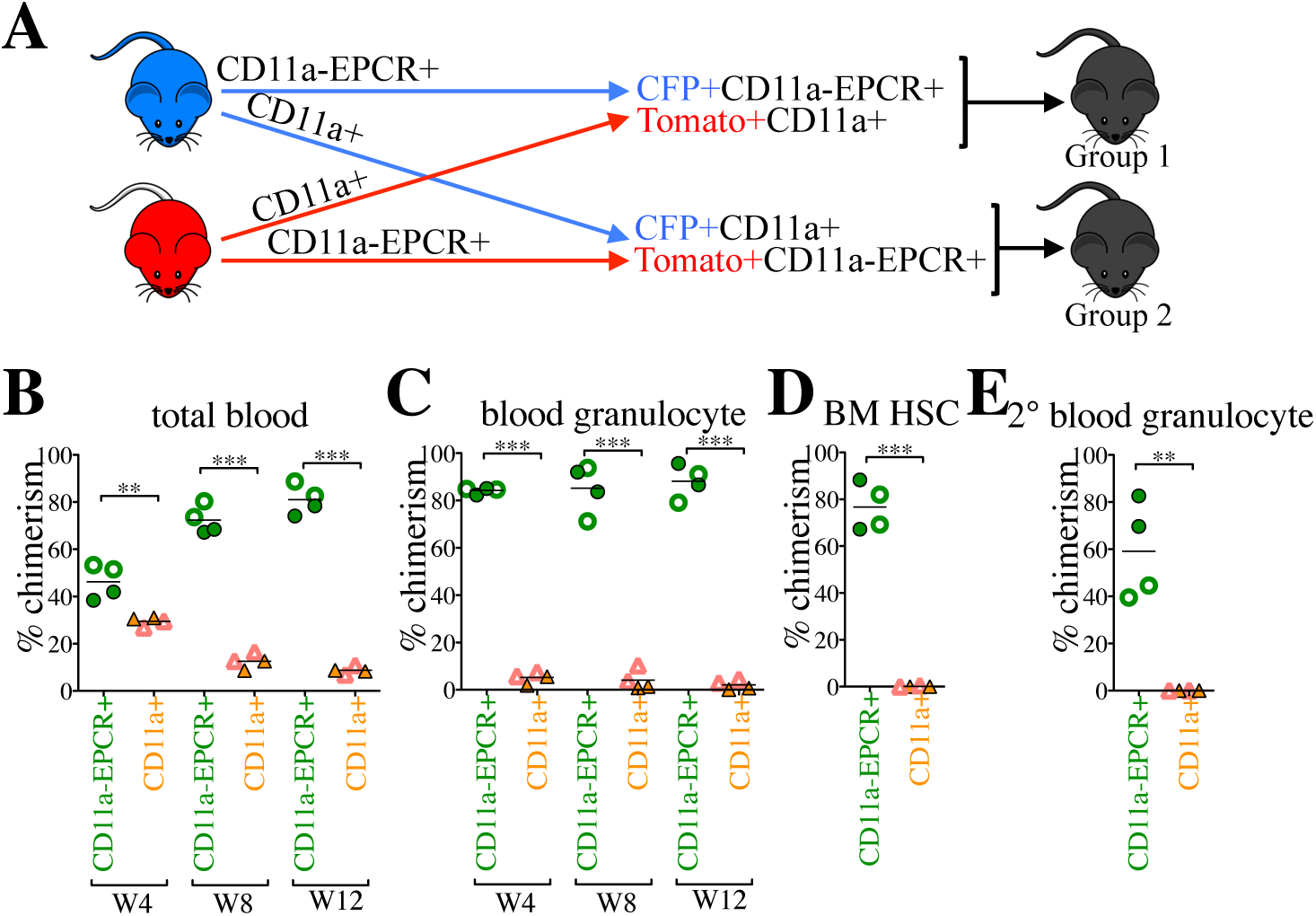
CD11a-EPCR+ KLS cells outcompete CD11a+ counterparts in competitive transplants. **A)** Representation of competitive transplant system. Group 1 recipients (filled symbols in B-E) received CFP+ CD11a- EPCR+ KLS and Tomato+ CD11a+ KLS, and Group 2 (hollow symbols in B-E) received Tomato+ CD11a- EPCR+ KLS and CFP+ CD11a+ KLS. 3,000 CD11a- EPCR+ KLS and 10,000 CD11a+ KLS sorted cells (physiological ratios) along with 100,000 helper WBM (from Wt Bl/6 mice) were co-transplanted into each lethally-irradiated Bl/6 recipient. **B-C)** Time-course analysis of donor chimerism in blood. Total (**B**) and granulocyte (**C**) blood chimerism from CD11a- EPCR+ KLS (“CD11a- EPCR+”) and CD11a+ KLS (“CD11a+”) sources in primary recipients at weeks (W) 4, 8, and 12 post-transplant. **D)** Donor chimerism of HSCs in the BM of primary recipients 13 weeks post-transplant. HSCs are defined as Ter119- CD27+ Sca-1+ Kit+ CD11a- EPCR+. **E)** Blood granulocyte chimerism in secondary recipients 6 weeks post-secondary transplant. Secondary transplants were done using 1x10^6^ WBM harvested from primary recipients 13 weeks after the primary transplantation. *\*\*p ≤ 0.01, ***p ≤ 0.001 (Student’s unpaired t test).*

We also examined the distribution of lineages derived from the two populations (**Supporting Information Fig. S2**). CD11a+ KLS-derived cells showed a lymphoid bias due to significantly higher production of B cells when compared to CD11a- EPCR+ KLS and non-transplanted controls. This lymphoid bias suggests the CD11a+ KLS fraction contained mostly progenitors that predominantly give rise to long-lived B lymphocytes. These results indicate that essentially all HSCs within the KLS population are CD11a- EPCR+.

### CD11a and EPCR alone can enrich HSCs without the need for other HSC markers

We next tested whether CD11a and EPCR alone were sufficient to sort HSCs in the absence of all other HSC markers. We used only these two markers for a competitive transplantation assay, and did not include any other HSC markers. We sorted CD11a- EPCR+ (“11a/EPCR”) cells into one tube, and all other live cells (referred to as “Not 11a/EPCR”) into another tube, then co-transplanted one population from CFP+ BM and the other population from Tomato+ BM into the same recipient mice. By sorting all cells outside of the CD11a- EPCR+ gate, we could ensure that any potential HSCs that fall outside of the CD11a- EPCR+ population would be transplanted in the “Not 11a/EPCR” fraction. We sorted and transplanted 500,000 BM cells per recipient, but maintained the physiological ratios of CD11a- EPCR+ cells (∼0.17% of whole BM; WBM) and Not 11a/EPCR (∼99.1% of WBM) (Fig. 3A). Thus, the transplanted cells are the equivalent of 500,000 WBM cells, with the CD11a- EPCR+ cells distinguishable from the rest of the BM cells by CFP or Tomato expression. In the recipient mice, we found that only the CD11a- EPCR+ donor source showed donor chimerism in primary and secondary recipients (Fig. 3B-C). We also examined the BM and found that donor HSPCs were only derived from the CD11a- EPCR+ source (Fig. 3D). These results indicate that all HSCs are present in the rare CD11a- EPCR+ fraction of BM, and that CD11a and EPCR together are sufficient to sort an enriched HSC population.

**Figure 3.**
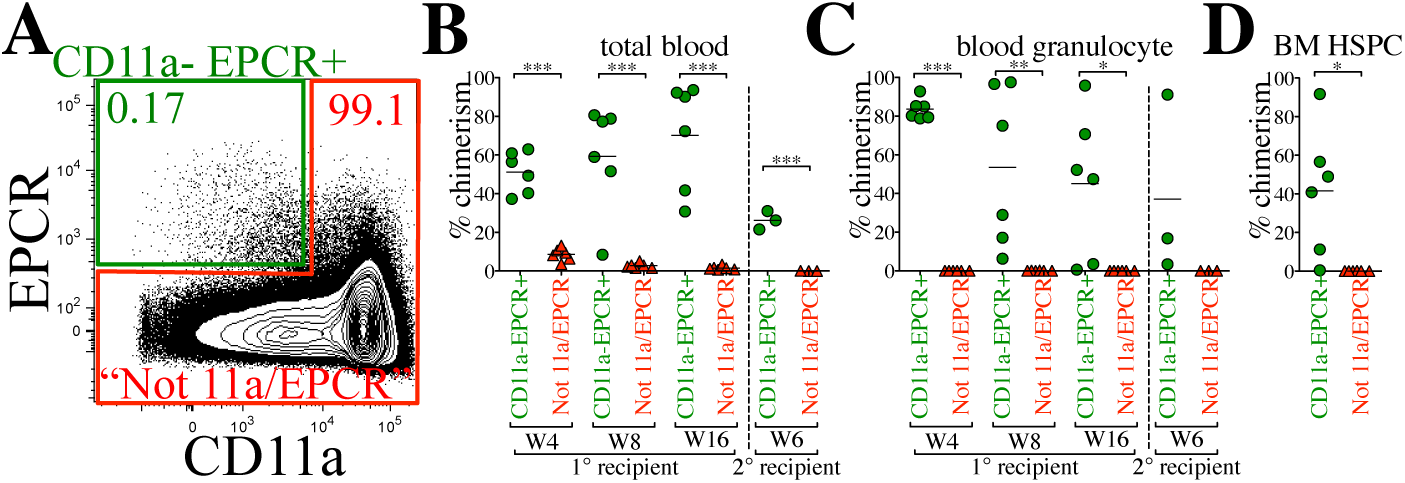
CD11a and EPCR alone are sufficient to sort rare population enriched for HSCs. **A)** Sorting strategy using only CD11a and EPCR as HSC markers. CFP+ CD11a- EPCR+ and Tomato+ “Not 11a/EPCR” (not CD11a- EPCR+) cells (and vice versa) were sorted and co-transplanted in a competitive setting and at the ratios shown. 850 CD11a- EPCR+ and 500,000 Not 11a/EPCR were transplanted into each recipient. Percentages of cells within each gate are shown. **B-C)** Time-course analysis of total blood **(B)** and blood granulocyte **(C)** chimerism from CD11a- EPCR+ and Not 11a/EPCR sources in primary
recipients 4, 8, and 16 weeks (W) post-transplant, and in secondary recipients (separated by vertical dashed line) at W6 following secondary transplant. Not all primary recipients were selected for secondary transplantation. **D)** Donor chimerism of HSPCs in the BM of primary recipients transplanted with “CD11a- EPCR+” and “Not 11a/EPCR” sorted cells 17 weeks post-transplant. HSPCs are defined as Ter119- CD27+ Sca-1+ Kit+. *\*p ≤ 0.05, **p ≤ 0.01, ***p ≤ 0.001 (Student’s unpaired t test). “Not 11a/EPCR”=not CD11a-EPCR+.*

### “11a/EPCR” two-color sorting method produces similar purity of HSCs as the “SLAM” method

While all HSCs are contained within the CD11a- EPCR+ fraction, it does not mean the population contains only HSCs. When examining BM cells gated on CD11a- EPCR+, approximately 81% are Kit+ and Sca-1+, but only 12% are CD150+ CD48- (**Supporting Information Fig. S3A**). It was previously shown that HSCs are CD150+ CD48-, and thus there are likely non-HSCs within the CD11a- EPCR+ fraction. The SLAM markers CD150 and CD48 have also been shown to be sufficient for two-color sorting of HSCs [6]. However, when examining the SLAM fraction (CD150+ CD48-) of BM, only 11% of these cells were Kit+ Sca-1+, and only 7.2% were CD11a- EPCR+. To confirm the efficiency of the SLAM method to purify HSCs in our hands, we transplanted CD150+ CD48- (SLAM) and “Not” CD150+ CD48- (referred to as “Not SLAM”) in a competitive setting, and found that only the SLAM cells were able to engraft long-term (**Supporting Information Fig. S3B-D**). Thus, it appears likely that all HSCs are contained within the SLAM population, but there are also many non-HSCs in this population.

To directly compare our “11a/EPCR” two-color sorting method with the SLAM method, CD11a- EPCR+ (11a/EPCR) and CD150+ CD48- (SLAM) cells were sorted, mixed, and co-transplanted into recipients in a competitive setting (Fig. 4A, Group 1 recipients). Equal numbers of each population (380 cells) were transplanted, allowing us to directly compare which population contained the most HSCs. While at week 4 the 11a/EPCR population had greater granulocyte chimerism than SLAM, after week 4 no statistical difference in granulocyte chimerism between the two populations was detected in primary and secondary recipients (Fig. 4Bi). Analysis of HSPCs in the BM also confirmed comparable engraftability between the two populations (Fig. 4Bii). Total blood chimerism and lineage distribution was not significantly different between the two sorting methods in primary and secondary recipients (**Supporting Information Fig. S4A-B**).

**Figure 4.**
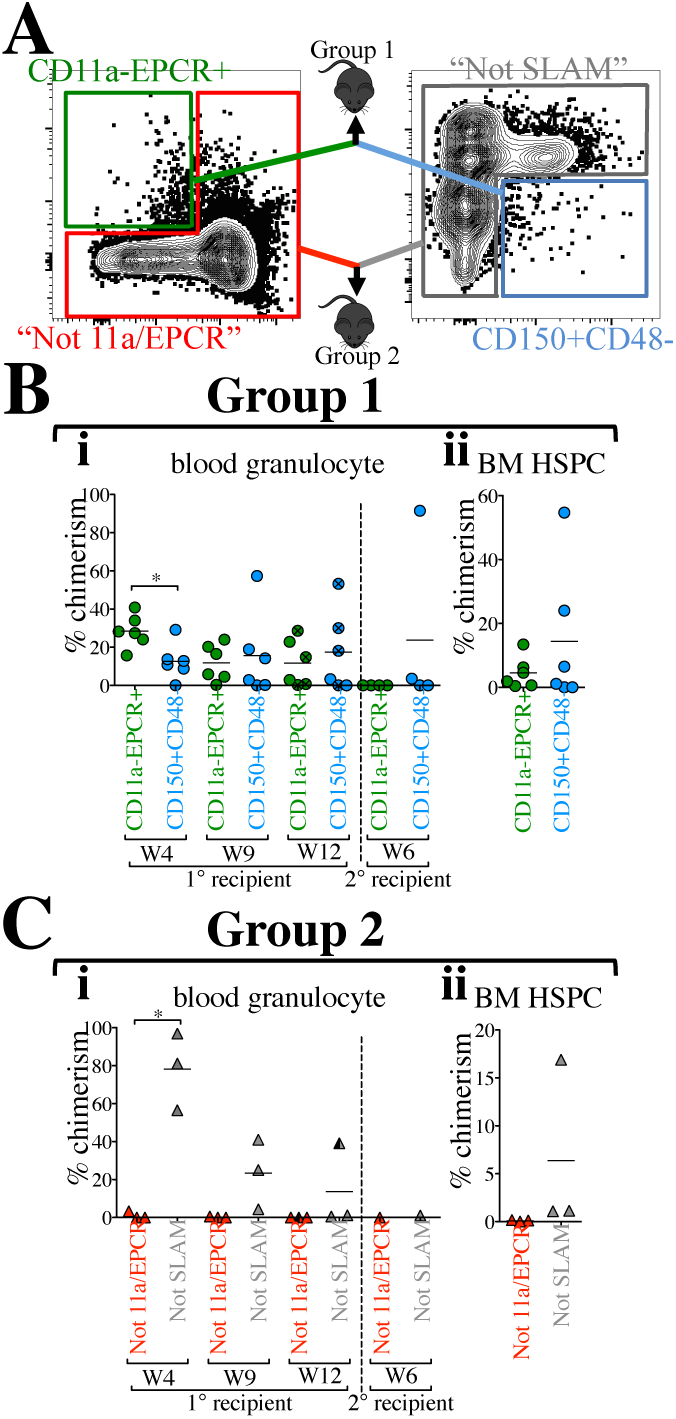
“11a/EPCR” two-color sorting is as efficient as using the “SLAM” method. **A)** Representation of direct comparison of two-color sorting methods. BM from CFP or Tomato mice was sorted using the combination of CD11a and EPCR only or the combination of CD150 and CD48 only. Approximately 380 cells from each of CD11a- EPCR+ and CD150+ CD48- gates were sorted, mixed, and co-transplanted with added 250,000 helper/competitor WBM (Group 1 recipients). Percentages of cells within each gate was kept consistent between the two methods. 200,000 cells from outside of the CD11a- EPCR+ gate (“Not 11a/EPCR”) and outside of the CD150+ CD48- gate (“Not SLAM”) were also mixed and co-transplanted (Group 2 recipients). **B)** i)Time-course analysis of blood granulocyte chimerism from CD11a- EPCR+ and CD150+ CD48- sources in primary recipients 4, 9, and 12 weeks (W) post-transplant, and in secondary recipients (separated by vertical dashed line) at week 6 following secondary transplant. Primary recipients used for secondary transplants are marked with an “x” inside circles at the 12-week timepoint. ii) Donor chimerism of HSPCs in the BM of primary recipients transplanted with CD11a- EPCR+ and CD150+ CD48- sorted cells 13 weeks post-transplant. **C)** i) Time-course analysis of blood granulocyte chimerism from “Not 11a/EPCR” and “Not SLAM” sources in primary recipients 4, 9, and 12 weeks (W) post-transplant, and in secondary recipients (separated by vertical dashed line) at week 6 following secondary transplant. The primary recipients used for secondary transplant is marked (half shaded black) at the 12-week timepoint. ii) Donor chimerism of HSPCs in the BM of primary recipients transplanted with “Not 11a/EPCR” and “Not SLAM” sorted cells 13 weeks post-transplant. The difference in chimerism is not statically significant. *\*p ≤ 0.05 (Student’s unpaired t test). “Not 11a/EPCR”= not CD11a-EPCR+; “Not SLAM”=not CD150+CD48-.*

To determine whether any HSCs resided outside of the CD11a- EPCR+ or the CD150+ CD48- gates, we sorted each of the “Not” populations (“Not 11a/EPCR” and “Not SLAM”) and co-transplanted them into recipient mice (Fig. 4A, Group 2 recipients). Even though we detected significantly higher “Not SLAM”-derived granulocytes compared to “Not 11a/EPCR” at week 4 post-transplant, at later timepoints and in secondary transplants, no significant difference was detected between the two methods (Fig. 4Ci). This indicates that no HSCs are found outside of the SLAM or 11a/EPCR fractions. However, we found high total chimerism and lymphocyte chimerism from the Not SLAM population, suggesting multipotent and lymphocyte progenitors were highly enriched in this population (**Supporting Information Fig. S4C**). Our data indicate that both two-color strategies are effective at sorting HSCs, although neither is pure for HSCs.

### CD11a and EPCR combination identifies phenotypic HSCs following irradiation, but not in inflammation

Many common HSC markers change their expression when challenged, such as during an inflammatory response. Thus, the phenotypic definition of HSCs can change depending on the context. We sought to determine the expression levels of CD11a and EPCR and their ability to mark HSCs after two types of challenge: irradiation and LPS-induced inflammation (Fig. 5). We sub-lethally irradiated (6 Gy) Bl/6 mice and examined their BM 48 hours post-irradiation. Consistent with previous observations, we detected a dramatic decrease in Kit expression[12, 14], reducing the frequency of KLS cells (Fig. 5A). The percentage of CD11a+ cells appeared to increase slightly after irradiation, though EPCR expression appeared unchanged (Fig. 5B, **Supporting Information Fig. S5A**). Other HSC markers also appeared unchanged, with the exception of CD48 which decreased after irradiation (**Supporting Information Fig. S5A**). Although CD11a expression appeared to increase overall in the irradiated BM, phenotypic HSCs were still found in the CD11a- EPCR+ fraction, suggesting these markers could still identify HSCs after irradiation (Fig. 5C, **Supporting Information Fig. S5B**).

**Figure 5.**
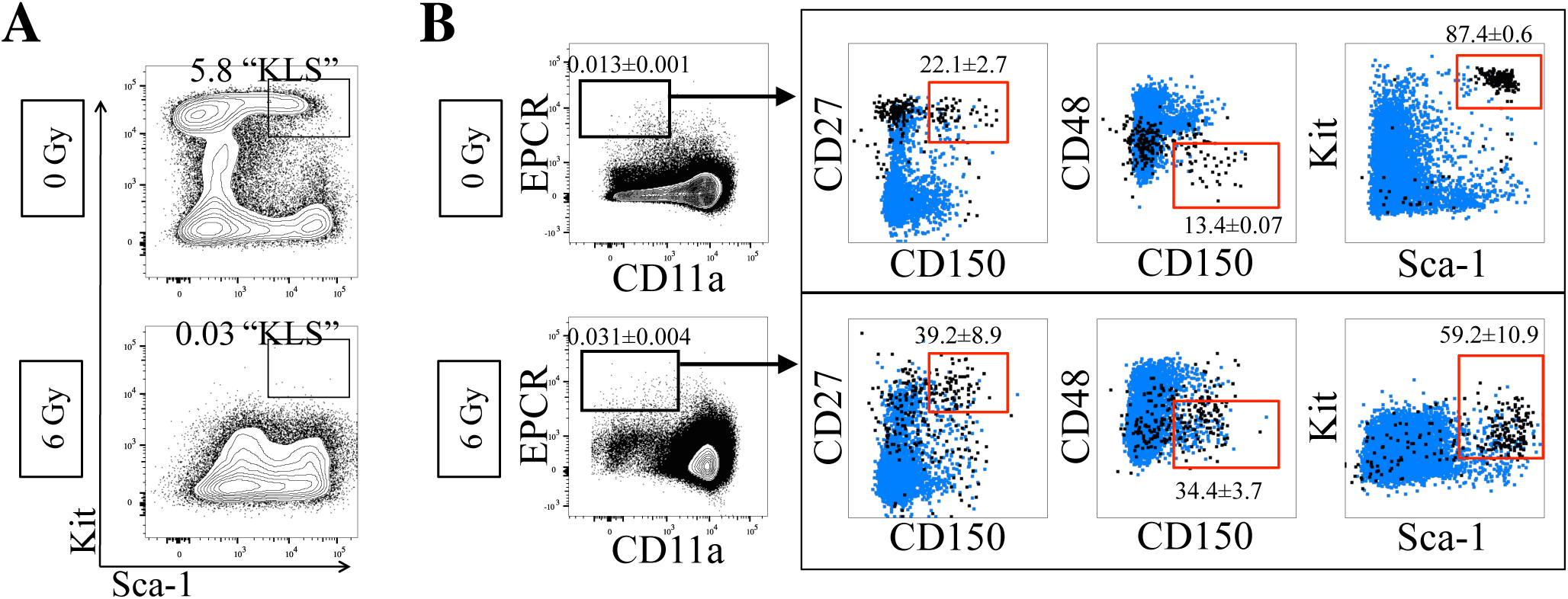
Efficacy of CD11a/EPCR combination post-irradiation injury. **A**) Expression of Sca-1 and Kit on Ter119- CD27+ BM cells in non-irradiated controls (top) compared to 48 hours after 6 Gy irradiation (bottom). Percentages of KLS cells are shown. **B**) Expression of HSC markers on Ter119- CD11a- EPCR+ without (0 Gy; n=3) and with (6 Gy; n=5) irradiation-induced BM injury. Numbers shown are percentages of cells within CD11a- EPCR+ gate ±SD. Boxed plots show Ter119- CD11a- EPCR+ gated cells (black) and total Ter119- cells (blue). Gates (red) show phenotypic HSCs and what percentage of Ter119- CD11a- EPCR+ cells fall within those gates.

We also examined CD11a and EPCR expression after LPS-induced inflammation, which is known to upregulate Sca-1 and therefore make HSC identification more difficult [24, 25]. While CD11a expression appeared unchanged in LPS-treated BM, EPCR expression changed significantly, and thus we were unable to use this marker combination to identify phenotypic HSCs in this context (**Supporting Information Fig. S6**). CD11a could still be combined with other HSC markers that appeared unchanged in inflammation, including CD27 and Kit. Future transplantation experiments would be required to confirm whether the phenotypic HSCs identified in either irradiation or inflammation are in fact functional HSCs.

## Discussion

Here, we demonstrate a novel strategy for a simplified, reproducible, and efficient way for HSC sorting with the use of CD11a and EPCR. Our transplantation strategy used direct competition between the two KLS fractions as the primary method to evaluate which fraction contained the most HSCs. Methods like limit dilution assays and single cell transplantation assays can provide quantitative estimates of the number of HSCs in a population. While we did not perform those types of assay, in our system we transplanted all possible sources of HSCs at their physiologic proportions into each recipient. Thus, if more HSCs existed outside of the CD11a- EPCR+ fraction, then the “Not CD11a- EPCR+” fraction would have shown higher donor chimerism. The fact that little granulocyte chimerism was found outside of the CD11a- EPCR+ fraction indicates that this population contained all HSCs.

We found that the 11a/EPCR method was comparable to the SLAM method of two-color HSC sorting. This is despite the fact that there appeared to be very little overlap between the two populations, with only 12% of CD11a- EPCR+ cells falling within the CD150+ CD48- gate and only 7.2% of CD150+ CD48- cells falling within the CD11a- EPCR+ gate. Because both methods have been shown to contain all HSCs, this means that likely all HSCs exist within the overlapping population, which would be CD11a- EPCR+ CD150+ CD48-. This also indicates that both populations contain many non-HSCs, as expected from a two-color approach. For the CD11a- EPCR+ fraction, the contaminating cells are highly enriched for MPPs, as nearly all the cells (81%) were Kit+ Sca-1+. For the SLAM fraction, most of the contaminating cells were CD11a+ and possibly lymphoid or myeloid cells. Therefore, both strategies have their strengths and weaknesses for use as two-color method, and the user should select whether they would rather have MPPs in their sort (11a/EPCR method) or other contaminating cells which are likely not progenitors (SLAM method).

Use of CD11a and EPCR to identify HSCs after irradiation or induction of inflammation gave mixed results. The combination appeared to work after irradiation, although total levels of CD11a were upregulated. CD11a upregulation may be involved in HSC differentiation to progenitors, which could be necessary to replenish hematopoietic populations depleted by irradiation, and thus the true undifferentiated HSCs remain CD11a-. As part of LFA-1, CD11a upregulation may also be involved in the migration of HSCs out of their niche and into circulation.

Efficient sorting of mouse HSCs allows in-depth molecular and functional characterization and contributes greatly to our understanding of the biology of these cells. However, the identification of human HSC markers promises a directly therapeutic impact. Recently, EPCR has been utilized for purification of *in vitro* expanded human HSCs [26]. Whether or not CD11a can similarly be used for human HSC identification is of great translation interest, and merits further examination.

## Acknowledgments

The authors wish to thank Martina Sassone-Corsi, Melissa Lodoen, Craig Walsh, and Matthew Blurton-Jones for technical assistance and helpful discussion, and Yasamine Ghorbanian, Ankita Shukla, and Tannaz Faal of the Inlay lab for helpful comments and discussion. This study was supported by National Institutes of Health grant R56HL133656 (to M.A.I.), American Cancer Society ACS/IRG Seed Grant IRG 98-27-10 (to M.A.I.), and California Institute for Regenerative Medicine grant CL1-00520-1.2 (to V.S. and the UC Irvine Stem Cell FACS Core).

The authors have no financial conflicts of interest to disclose.

## References

1. Seita J, Weissman IL. Hematopoietic stem cell: self-renewal versus differentiation. Wiley interdisciplinary reviews Systems biology and medicine. 2010;2:640–653.

2. de la Morena MT, Nelson RP, Jr. Recent advances in transplantation for primary immune deficiency diseases: a comprehensive review. Clinical reviews in allergy & immunology. 2014;46:131–144.

3. Boitano AE, Wang J, Romeo R et al. Aryl hydrocarbon receptor antagonists promote the expansion of human hematopoietic stem cells. Science. 2010;329:1345–1348.

4. Brunstein CG, Gutman JA, Weisdorf DJ et al. Allogeneic hematopoietic cell transplantation for hematologic malignancy: relative risks and benefits of double umbilical cord blood. Blood. 2010;116:4693–4699.

5. Hagedorn EJ, Durand EM, Fast EM et al. Getting more for your marrow: boosting hematopoietic stem cell numbers with PGE2. Experimental cell research. 2014;329:220–226.

6. Kiel MJ, Yilmaz OH, Iwashita T et al. SLAM family receptors distinguish hematopoietic stem and progenitor cells and reveal endothelial niches for stem cells. Cell. 2005;121:1109–1121.

7. Balazs AB, Fabian AJ, Esmon CT et al. Endothelial protein C receptor (CD201) explicitly identifies hematopoietic stem cells in murine bone marrow. Blood. 2006;107:2317–2321.

8. Karlsson G, Rorby E, Pina C et al. The tetraspanin CD9 affords high-purity capture of all murine hematopoietic stem cells. Cell reports. 2013;4:642–648.

9. Wiesmann A, Phillips RL, Mojica M et al. Expression of CD27 on murine hematopoietic stem and progenitor cells. Immunity. 2000;12:193–199.

10. Chen JY, Miyanishi M, Wang SK et al. Hoxb5 marks long-term haematopoietic stem cells and reveals a homogenous perivascular niche. Nature. 2016;530:223–227.

11. Challen GA, Boles N, Lin KK et al. Mouse hematopoietic stem cell identification and analysis. Cytometry Part A: the journal of the International Society for Analytical Cytology. 2009;75:14–24.

12. Simonnet AJ, Nehme J, Vaigot P et al. Phenotypic and functional changes induced in hematopoietic stem/progenitor cells after gamma-ray radiation exposure. Stem cells. 2009;27:1400–1409.

13. Baldridge MT, King KY, Goodell MA. Inflammatory signals regulate hematopoietic stem cells. Trends in immunology. 2011;32:57–65.

14. Vazquez SE, Inlay MA, Serwold T. CD201 and CD27 identify hematopoietic stem and progenitor cells across multiple murine strains independently of Kit and Sca-1. Experimental hematology. 2015;43:578–585.

15. Kinashi T. Intracellular signalling controlling integrin activation in lymphocytes. Nature reviews Immunology. 2005;5:546–559.

16. Zhang Y, Wang H. Integrin signalling and function in immune cells. Immunology. 2012;135:268–275.

17. Lum AF, Green CE, Lee GR et al. Dynamic regulation of LFA-1 activation and neutrophil arrest on intercellular adhesion molecule 1 (ICAM-1) in shear flow. The Journal of biological chemistry. 2002;277:20660–20670.

18. Shamri R, Grabovsky V, Gauguet JM et al. Lymphocyte arrest requires instantaneous induction of an extended LFA-1 conformation mediated by endothelium-bound chemokines. Nature immunology. 2005;6:497–506.

19. Fathman JW, Fernhoff NB, Seita J et al. Upregulation of CD11A on hematopoietic stem cells denotes the loss of long-term reconstitution potential. Stem cell reports. 2014;3:707–715.

20. Benz C, Copley MR, Kent DG et al. Hematopoietic stem cell subtypes expand differentially during development and display distinct lymphopoietic programs. Cell stem cell. 2012;10:273–283.

21. Iwasaki H, Arai F, Kubota Y et al. Endothelial protein C receptor-expressing hematopoietic stem cells reside in the perisinusoidal niche in fetal liver. Blood. 2010;116:544–553.

22. Serwold T, Ehrlich LI, Weissman IL. Reductive isolation from bone marrow and blood implicates common lymphoid progenitors as the major source of thymopoiesis. Blood. 2009;113:807–815.

23. Inlay MA, Bhattacharya D, Sahoo D et al. Ly6d marks the earliest stage of B-cell specification and identifies the branchpoint between B-cell and T-cell development. Genes & development. 2009;23:2376–2381.

24. Zhang P, Nelson S, Bagby GJ et al. The lineage-c-Kit+Sca-1+ cell response to Escherichia coli bacteremia in Balb/c mice. Stem cells. 2008;26:1778–1786.

25. Esplin BL, Shimazu T, Welner RS et al. Chronic exposure to a TLR ligand injures hematopoietic stem cells. Journal of immunology. 2011;186:5367–5375.

26. Fares I, Chagraoui J, Lehnertz B et al. EPCR expression marks UM171-expanded CD34+ cord blood stem cells. Blood. 2017;129:3344–3351.

